# Identification of host-interacting domains of surface layer protein B from *Levilactobacillus brevis* and the evidence as glycoprotein

**DOI:** 10.1101/2023.04.06.534471

**Authors:** Zheng Lin Tan, Naoyuki Yamamoto

**Affiliations:** School of Life Science and Technology, Tokyo Institute of Technology, Yokohama, 226-8501, Japan

**Keywords:** Levilactobacillus brevis, Surface layer protein, Glycoprotein, Dendritic cells, C-type lectins

## Abstract

Surface layer proteins (Slps) can be found on the surface of some lactobacilli which involves in host-interaction and show immunomodulatory activity. Despite the capability of Slps to interact with C-type lectins on antigen presenting cells, present of sugar has not been confirmed except for *Lentilactobacillus kefiri*. In this study, we investigated the presence of sugar moiety in the structure of SlpB from *Levilactobacillus brevis*, and its involvement in interaction with DC-SIGN of dendritic cells. 4 fragments of sugar containing fragments were identified in SlpB, and the involvement of sugar in DC-SIGN interaction was confirmed by competitive assay against hexoses. The sequences of these fragments were identified based on the mass obtained from mass spectrometry, and the result has confirmed that these fragments were from carbohydrate binding motifs of SlpB. Analysing the binding affinities of these fragments with surface resonance plasmon analysis suggested that 2 fragments possess higher binding affinity compared to the remaining fragments, and the sum of binding affinities of these fragments was approximate the affinity of intact SlpB.

## Introduction

Lactobacilli are known to be abundant colonizer of human small intestinal mucosa which involves in host immunomodulation. Among them, *Levilactobacillus brevis (L. brevis)* JCM 1059 which is human origin, is recognized for its host interacting properties and induction of adjuvant effect through its surface layer protein (Slp) (Prado Acosta et al., 2021). When Slp was purified from the surface of *L. brevis* and coated on the surface of liposome, it could facilitate specific uptake into Peyer’s patch via microfold cells, and target antigen presenting cells lie in subepithelial dome (Tan et al., 2022). These results suggested that Slp from *L. brevis* plays an important role in host interaction and immunomodulation. However, detailed mechanisms of interaction of Slp to host remained unknown. To understand how *L. brevis* interact with host, particularly in host-microbial interplays, function of probiotic bacteria in intestine and to further extend the application of Slp from *L. brevis*, it is important to characterize the properties of Slp from *L. brevis* and elucidate its mechanism of action.

Professional antigen presenting cells, which includes both macrophage and dendritic cells (DCs) regularly interact with bacteria and involve in immune sampling. Particularly, immature DCs could migrate through bloodstream and actively interact with bacteria in the body. Then, then antigen were presented to T cells, which will result in differentiation of naïve T cells toward T helper (Th) 1, Th2, or T regulatory cells (Mohamadzadeh et al., 2005; Smits et al., 2005). Among the receptors expressed on the surface of antigen-presenting cells, particularly macrophage and DCs, C-type lectins are the dominant receptors known to interact with bacteria. Dendritic cell-specific intercellular adhesion molecule 3-grabbing nonintegrin (DC-SIGN) is a DCs specific C-type lectins, which could recognize mainly mannose and fucose (Valverde et al., 2020). Therefore, DC-SIGN is important in inducing various responses mediated by DCs. Several pathogens (Geijtenbeek et al., 2003; Koppel et al., 2005) and lactobacilli (Smits et al., 2005), and Slp from lactobacilli was found to be the key component which bind to DC-SIGN (Konstantinov et al., 2008; Prado Acosta et al., 2016).

Although it is generally regarded that Slps from *L. brevis* is non-glycosylated (Prado Acosta et al., 2016), it contradict with the property of C-type lectins, e.g., Mincle from macrophage (Feinberg et al., 2013) and DC-SIGN from DCs (Guo et al., 2004), which the interaction is mediated through sugar moiety at carbohydrate binding domain. The detailed molecular mechanism of binding of Slp, particularly Slp from *L. brevis* to DC-SIGN which included the binding domain of Slp, and whether protein modification exist on Slp from *L. brevis* and their involvement (if they exist) in DC-SIGN interaction, remained unclear. In this study, we investigated the ligand-receptor interaction between SlpB from *L. brevis* and DCs and the potential of immunomodulating effect of SlpB from *L. brevis*.

## Results

### Stimulation of DC with SlpB from *L. brevis*

Previous studies have suggested that Slps from lactobacilli could induce cytokines production by antigen presenting cells as measured by the production of cytokines (Konstantinov et al., 2008; Malamud et al., 2020; Prado Acosta et al., 2021). To determine immunomodulating effect of SlpB on DCs, DCs were co-incubated with either ovalbumin (OVA) or lipopolysaccharide (LPS) with or without SlpB for 24 h in 37°C, 5% CO_2_ humidified incubator. SlpB has enhanced Th1, Th2 and Th17 cytokines production by DCs after stimulation with OVA and LPS (Figure 1), which suggested that SlpB might act as an adjuvant to enhance immune response resulted from OVA and LPS stimulation.

**Figure 1.**
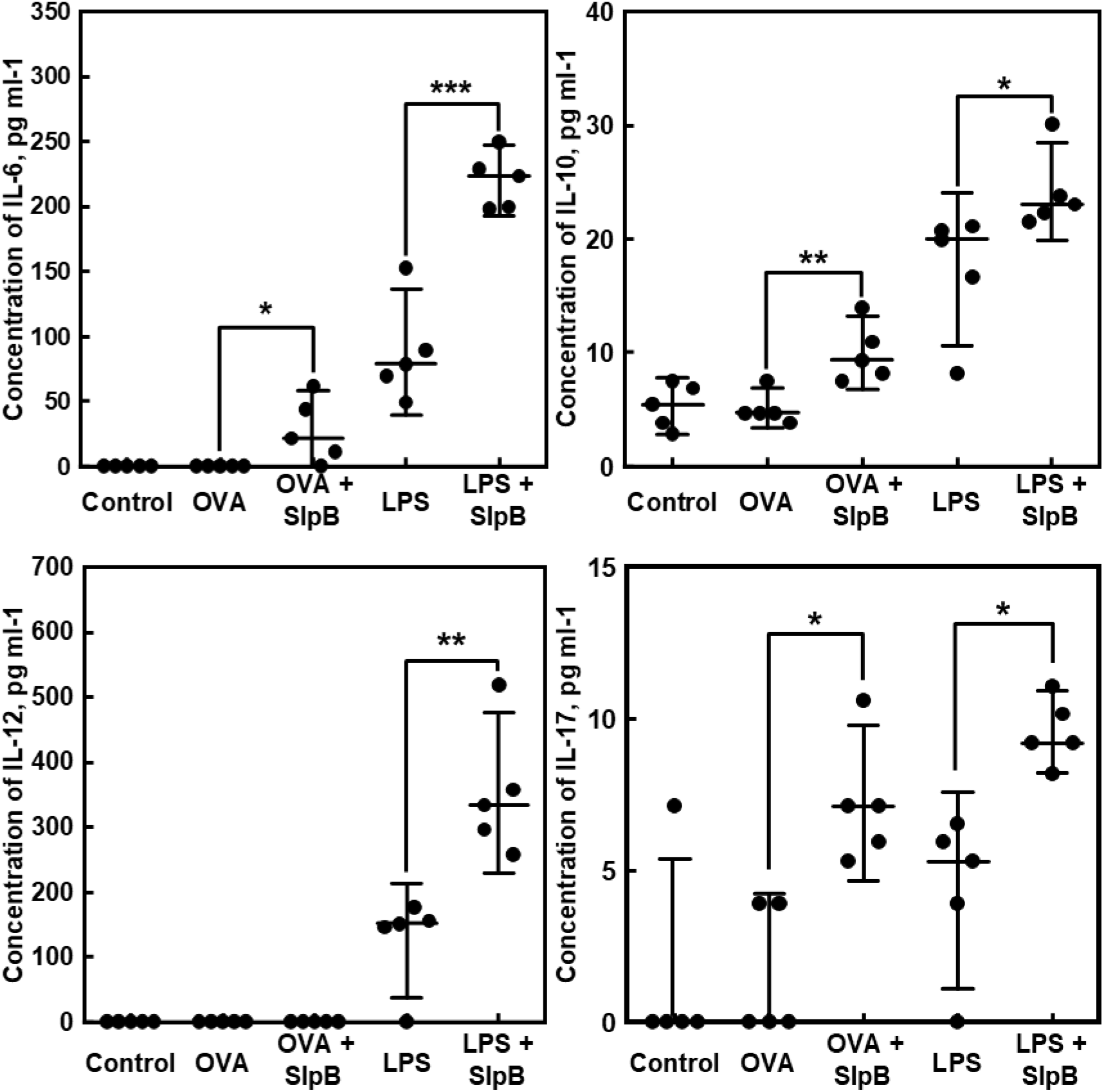
SlpB elicit cytokine responses in dendritic cells. The production of IL-6, IL-10, IL-12 and IL-17 by DC were analysed by ELISA. *n* = 5, median ± 95% confidence interval of mean. Statistical significance were evaluated with Student’s t-test. * *p* < 0.05, ** *p* < 0.01, *** *p* < 0.001.

### SlpB from *L. brevis* binds to DC-SIGN

Considering that it has been previously shown that SlpA from *Lactobacillus acidophilus* could binds to DC-SIGN of DCs (Konstantinov et al., 2008), we decided to test the ability of SlpB binding to DC-SIGN. Binding of SlpB to DCs was reduced after blocking DC-SIGN with antibody against DC-SIGN (Figure 2), which suggests that DC-SIGN is a receptor for SlpB from *L. brevis*.

**Figure 2.**
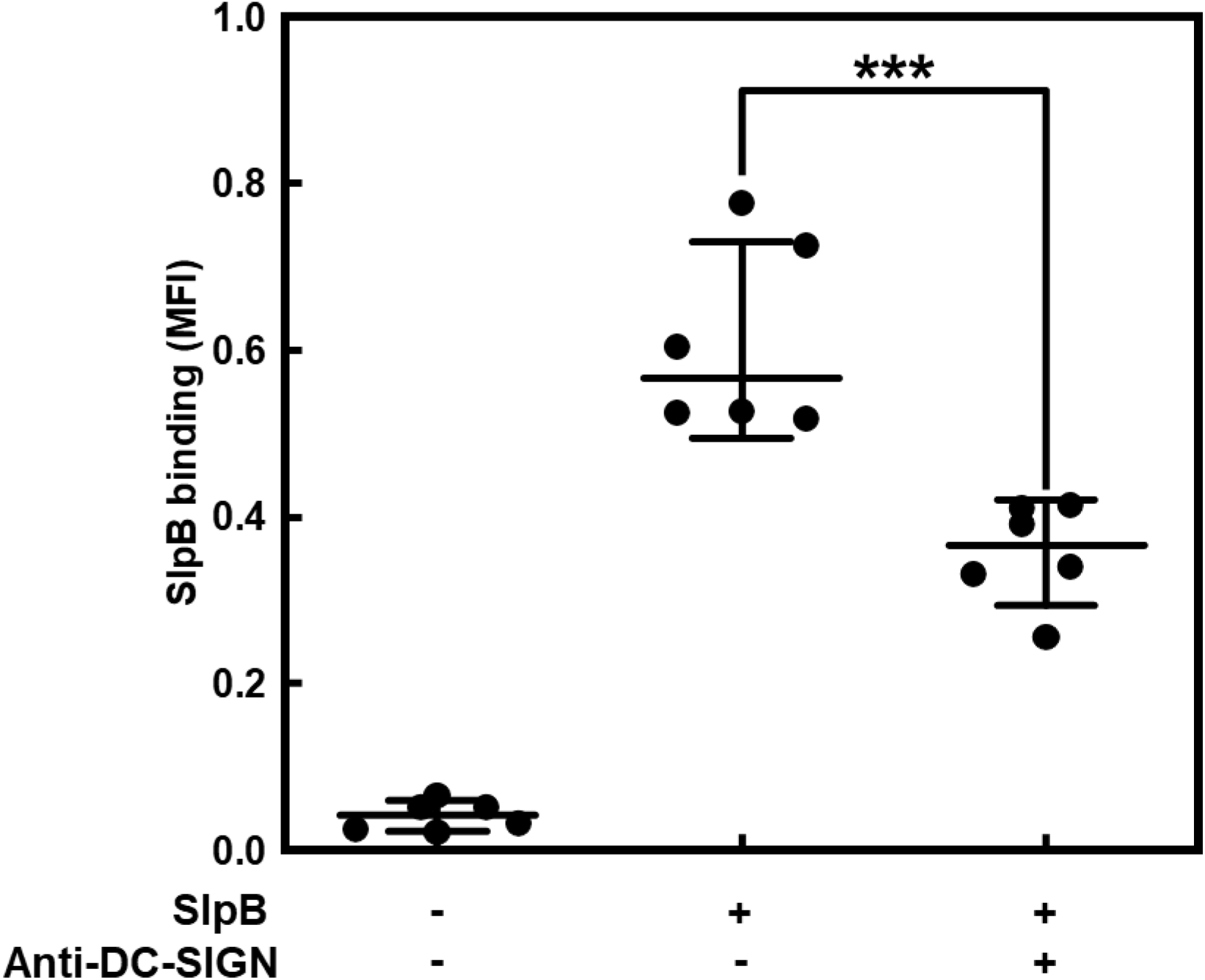
Cellular DC-SIGN is receptors for SlpB from *L. brevis*. *n* = 6, median ± 95% confidence interval of mean. Statistical significance were evaluated with Student’s t-test. \*\*\**p* < 0.001.

### SlpB from *L. brevis* is glycosylated and the involvement of sugar in DC interactions

To interact with C-type lectins, SlpB from *L. brevis* must contain carbohydrate chain (Guo et al., 2004; Valverde et al., 2020). However, prior studies have shown that SlpB is non-glycosylated through zymography (Prado Acosta et al., 2016). We decided to change an approach to investigate whether SlpB from *L. brevis* is glycosylated. Existence of carbohydrates in SlpB was confirmed with phenol-sulphuric acid method (Figure 3a). Then, SlpB was treated with PNGase F, and the presence of *N*-glycan was confirmed by sodium dodecyl sulphate polyacrylamide gel electrophoresis (SDS-PAGE) through the reduction of molecular size of SlpB after removal of *N*-glycan. A band with apparent molecular size of 48 kDa, which is in align with the molecular size of amino acid of SlpB appear after PNGase F treatment, suggested that SlpB contains *N*-glycan (Figure 3b).

**Figure 3.**
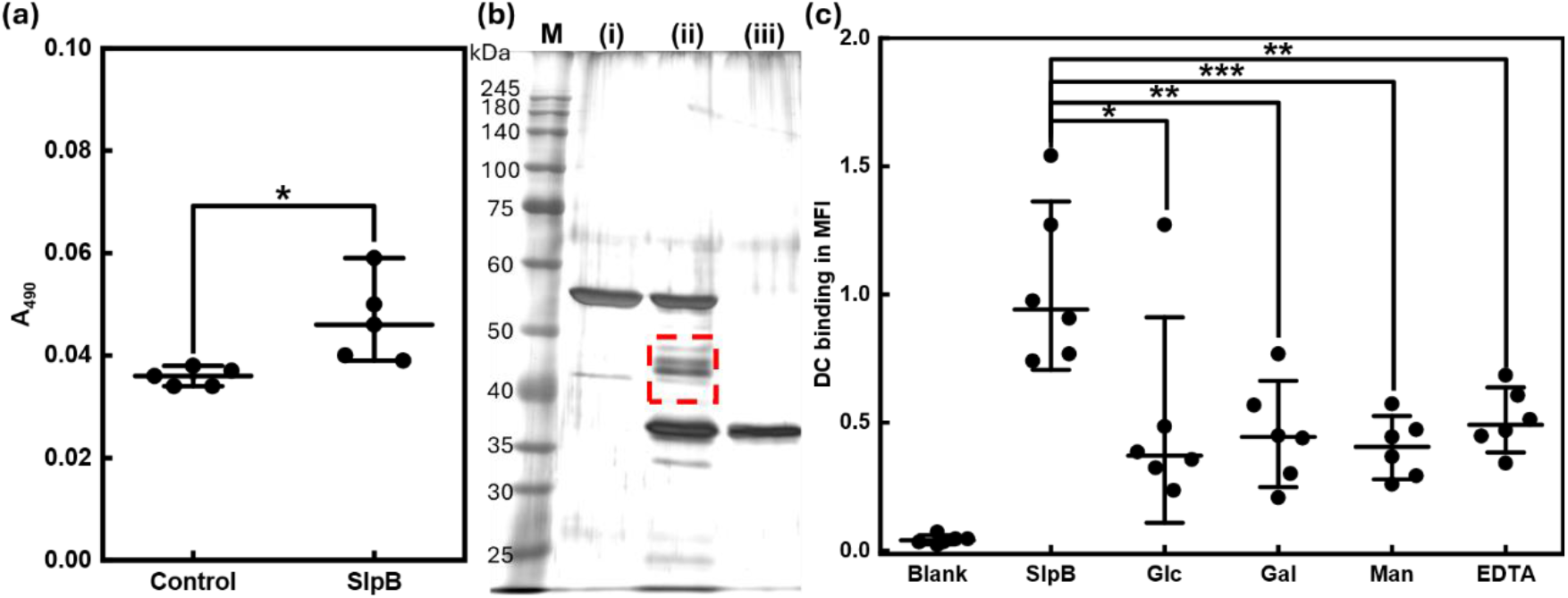
SlpB interacts with DC via sugar moiety. (a) Identification of carbohydrates in SlpB via phenol-sulphuric acid method. *n* = 5, median ± 95% confidence interval of mean. (b) SDS-PAGE of deglycosylated SlpB. (i) SlpB, (ii) SlpB co-incubated with PNGase F, (iii) PNGase F. Deglycosylated SlpB was gated with dashed line. (c) Competitive binding of SlpB to hexose. *n* = 6, median ± 95% confidence interval of mean. Statistical significances were evaluated with Student’s t-test. * *p* < 0.05, ** *p* < 0.01, *** *p* < 0.001.

To investigate the function of sugars which contain in SlpB in SlpB-DC interaction, we have conducted competitive binding assay of SlpB with hexoses, i.e., D-glucose (Glc), D-galactose (Gal), D-mannose (Man) and ethylenediaminetetraacetic acid (EDTA). Figure 3c shows both hexoses and EDTA could inhibit binding of SlpB to DC, which suggested that sugars in SlpB involved in DC interaction.

### Binding domains of SlpB to DC

To this point, we have identified *N*-glycan in SlpB, and shown that SlpB could binds to DC through sugar chain. Next, we would like to identify the domains of SlpB which involve in DC-interaction. Trypsinized SlpB was co-incubated with formalin-fixed DC for 60 min, and both trypsinized SlpB with (Figure 4a) or without (Figure 4b) incubation with DC was analysed with reversed phase high performance liquid chromatography (RP-HPLC). 4 peaks were attenuated after co-incubation with DC, which suggested that these peaks might contain fragment of SlpB which binds to DC. These fragments, together with another peak which did not attenuate (peak 5) was stained with Cy5, and co-incubated with DC to confirm the result obtained with RP-HPLC. Figure 4c shows fragments contained in peaks 1 – 4 bind to DC, while peak 5 did not bind to DC. This observation confirms the result obtained from RP-HPLC.

**Figure 4.**
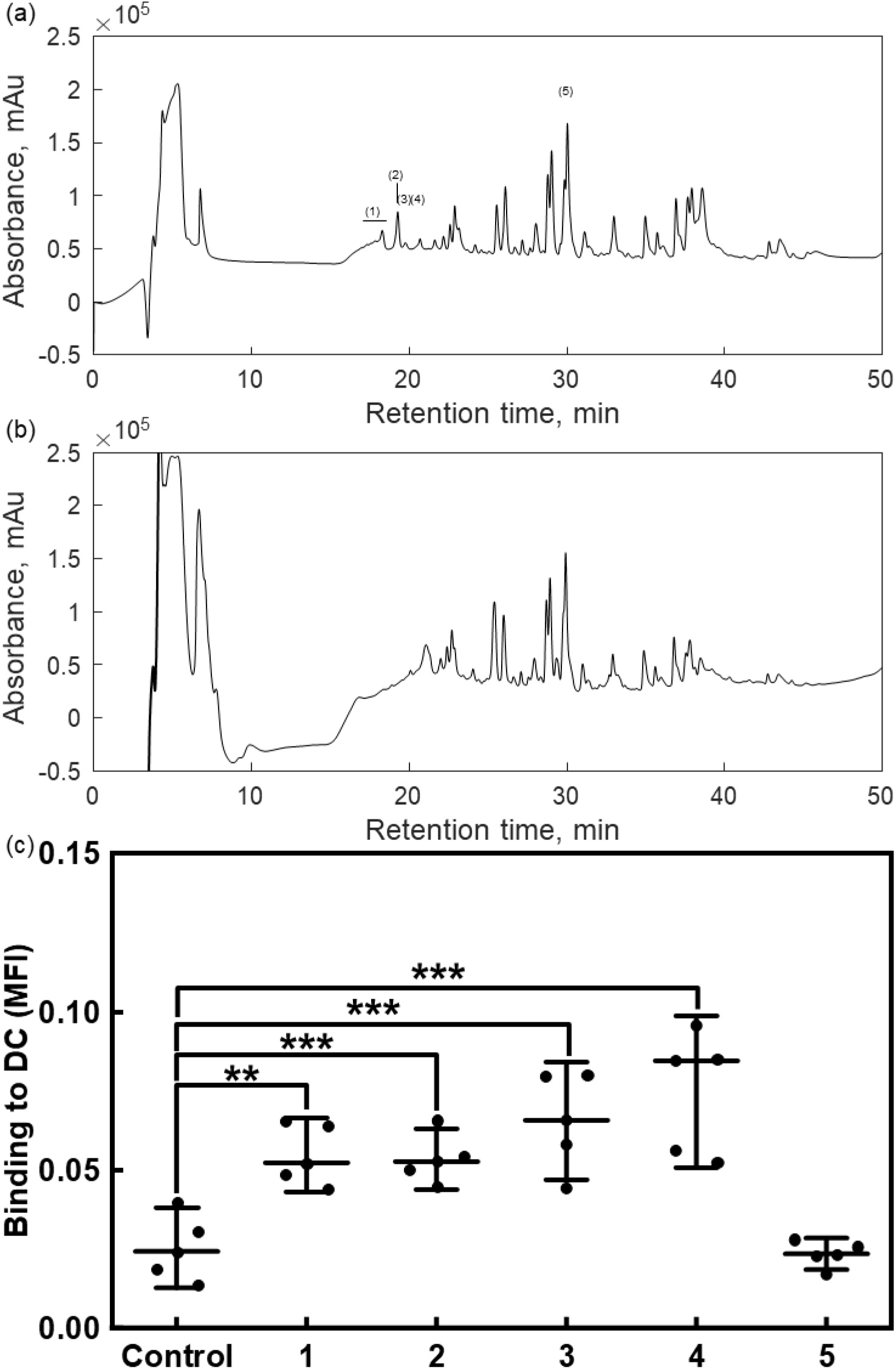
Fragments of SlpB responsible for binding to DC. Chromatogram of trypsinized SlpB (a) before and (b) after incubation with dendritic cells. (c) Binding of fragments of SlpB to DC. *n* = 5, median ± 95% confidence interval of mean. The number indicates the peak indicated in (a). Statistical significances were evaluated with Student’s t-test. * *p* < 0.05, ** *p* < 0.01, *** *p* < 0.001.

### Sugars involved in interaction of fragments of SlpB to DC

Since sugars involved in binding of SlpB to DC, and glycosylated fragments are usually highly hydrophilic, which will result in short retention time, these fragments of SlpB which bind to DC might be glycosylated domains of SlpB. To investigated whether these fragments bind to DC through sugars, competitive assays against Glc, Gal and Man were evaluated (Figure 5). The results suggested that hexoses inhibit the binding of these fragments to DC, with the Man exhibiting the strongest inhibitory activity, followed by Gal and Glc. Furthermore, the binding capability of fragments in peak 4 was the strongest (Figure 5d), followed by fragments in peak 3 > 2 > 1, measured by mean fluorescence intensity (MFI).

**Figure 5.**
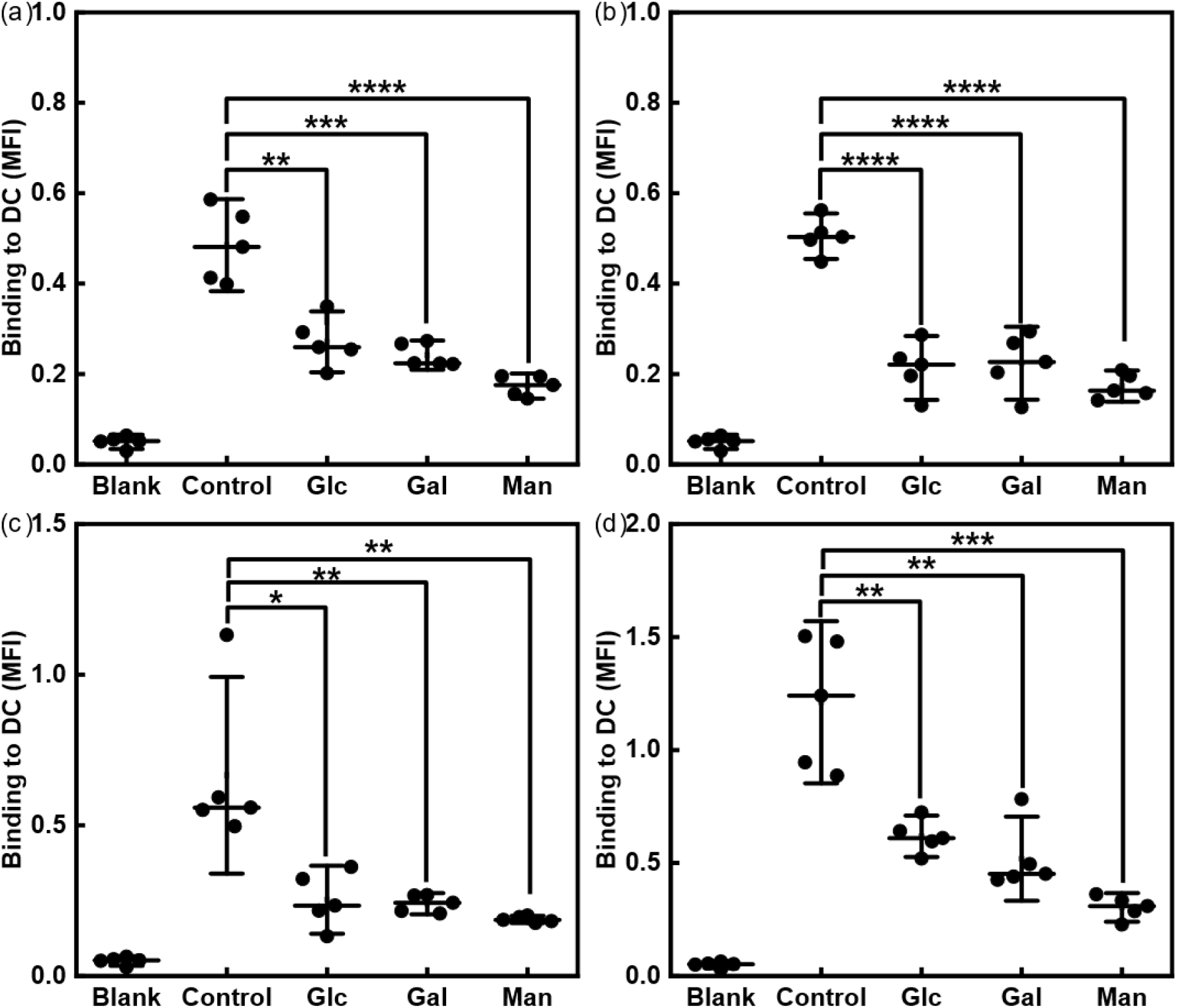
Fragments of SlpB interacts with DC via sugar moiety. Competitive binding assay of (a) fragment 1, (b) fragment 2, (c) fragment 3 and (d) fragment 4 with various hexose. *n* = 5, median ± 95% confidence interval of mean. Statistical significances were evaluated with Student’s t-test. * *p* < 0.05, ** *p* < 0.01, *** *p* < 0.001, **** *p* < 0.0001.

### Prediction of the sequences of peptide fragments

Given that the concentration of these peptides was low, it is difficult to conduct mass spectrometry/mass spectrometry (MS/MS) analysis or sequencing to identify the sequence of peptides. To predict the sequence of peptides, we first eliminate the non-binding peptides from the sequence of SlpB. The non-binding fragments of trypsinized SlpB were analysed with matrix assisted laser desorption/ionization time of flight-MS/MS (MALDI-ToF-MS/MS) to identify the sequence of non-binding fragments. Then, the sequences of non-binding fragments were eliminated from the sequence of SlpB, and the remaining fragments were the potential binding fragments (Figure 6).

**Figure 6.**
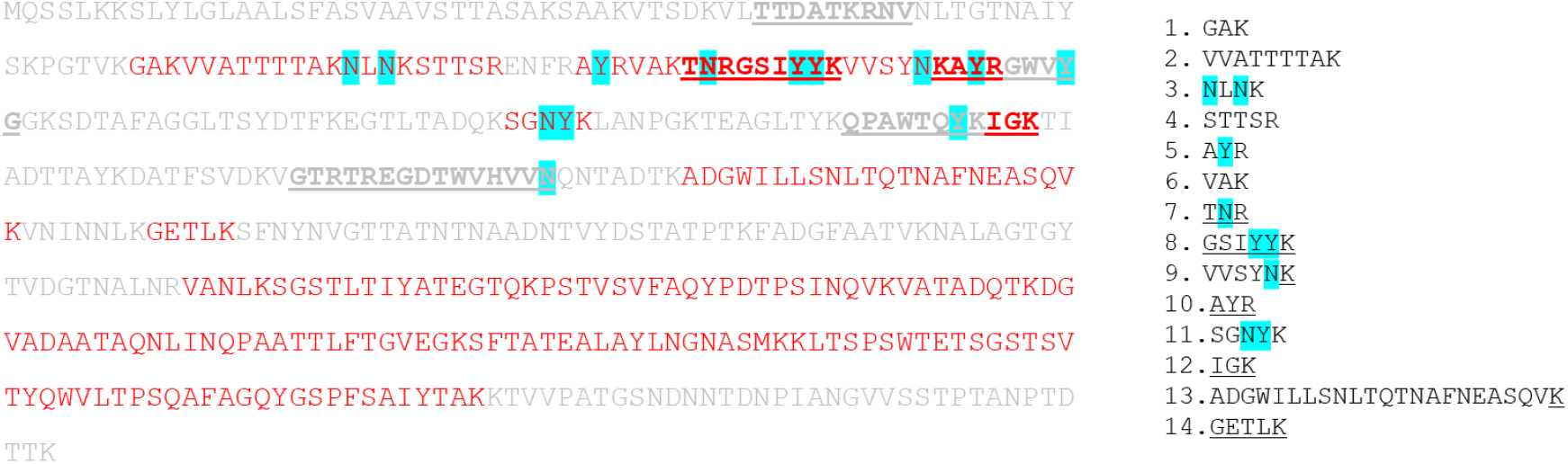
Potential fragments of SlpB which bind to DC. Fragments labelled in red represents fragments which bind to DC, Potential sugar binding amino acids are highlighted in cyan. Fragments at right indicates trypsinized fragments obtained from the fragments in red.

Based on the distribution of hydrophobic amino acid of SlpB and carbohydrates binding motifs (underlined in Figure 6), we observed that N-terminal of SlpB is hydrophilic and contains carbohydrate binding motifs, which make the sequences in N-terminal the candidate of the fragments in the 4 peaks. Then, *in silico* analysis of trypsin digestion of the candidate fragments in N-terminal was performed to identify the potential sequences located in N-terminal. After that, the fragments which bind to DC were collected, and digested with dipeptidyl peptidase 4 (DPP-IV) and PNGase F before MALDI ToF-MS analysis. The mass spectra suggest that the sequences of peptides in peak 1 – 4 were NLNK, AYR, SGNYK and GSIYYK respectively.

### Binding affinity of SlpB and its fragments to DC-SIGN

The binding affinity of SlpB and fragments 1 – 4 from SlpB to DC-SIGN were evaluated with Biacore system. Based on the sensogram, the dissociation constant, *kD* was calculated (Figure 7). *kD* value for fragments 1 – 4 and SlpB were 3.7 × 10^-5^, 4.07 × 10^-5^, 7.26 × 10^-7^, 1.95 × 10^-7^, 3.03 × 10^-8^ M respectively.

**Figure 7.**
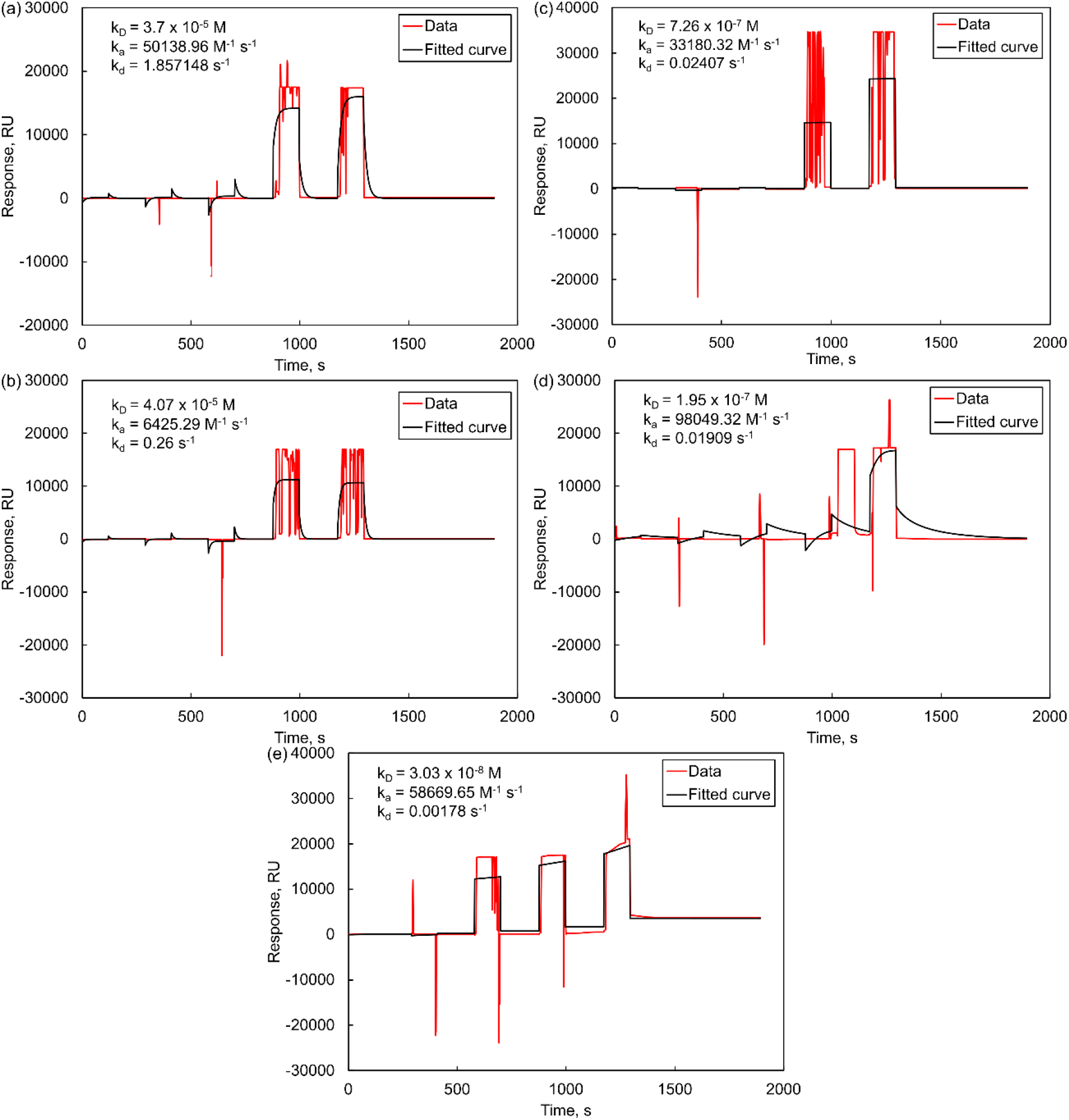
Affinity determination of SlpB and its fragments to DC-SIGN. Single cycle kinetics analysis was performed on DC-SIGN. Sensogram expressed in resonance unit for (a) Fragment 1, (b) fragment 2, (c) fragment 3, (d) fragment 4, (e) SlpB.

## Discussion

Glycosylation is a post-translational modification of proteins which has widely been found in surface-exposed proteins, e.g., flagellin and pilin. Although Slps expressed in various lactobacilli, e.g., *Lentilactobacillus kefiri* (Mobili et al., 2009), *Lentilactobacillus buchneri* (Möschl et al., 1993) and *Lactobacillus acidophilus* (Fina Martin et al., 2019), little was known about other genera of lactobacilli. Glycosylation of proteins involves in immunomodulation (Malamud et al., 2020), in addition to host interaction. In this study, we have shown that SlpB from *L. brevis* JCM 1059 is glycosylated (Figure 3). Carbohydrate chains in SlpB involves in interaction with DC (Figure 3), potentially through DC-SIGN binding. Furthermore, a large portion of glycosylated SlpB remained after co-incubation with PNGase F, suggested that the structure of *N*-glycan of SlpB is stable against enzymatic hydrolysis of *N*-glycosidase.

Trypsinization of SlpB, and analysis of the fragments obtained suggested that 4 fragments of peptides obtained from SlpB involved in DC interaction (Figure 4). Competitive assay with hexoses suggested that these fragments contain *N*-glycan, which the interaction with DC can be inhibited by hexoses (Figure 5). Considering the concentration of these peptides measured in absorbance at 220 nm (Figure 4), and the binding to DC (Figure 5), the strength of binding of these fragments to DC was 4 > 3 > 2 ≈ 1, which the affinity was in agree with the measurement obtained from Biacore analysis (Figure 7), while the inhibitory activity of hexose to the binding of these peptides to DC was Man > Gal > Glc.

After hydrolysis with both DPP-IV and PNGase F, increase in signal intensity in mass spectra which resemble the peptide fragments of the candidate of binding fragments and the dipeptides resulted from enzymatic hydrolysis by DPP-IV were identified. All fragments contain either N or Y, which might be *N*-glycosylated. Interestingly, these fragments were originated predominantly from 2 carbohydrates binding motifs from SlpB. The remaining carbohydrate binding motifs which potentially exhibit lectins-like activity (underlined, in grey in Figure 6), i.e., ^43^TTDATKRNV_51_ and ^197^GTRTREGDTWVHVVN_211_ were successfully analysed and assigned through y-ion by MALDI-ToF-MS/MS analysis from non-binding fragments, which suggest that these domains might be the cell wall binding domain of SlpB, rather than host-interacting domains. However, further analysis on the structure of glycans and the sequence of these 4 fragments isolated is required to reveal the reason which results in the difference in binding affinity.

As glycosylation is involved in immunomodulation of bacterial components, e.g., LPS, the adjuvant effect of SlpB which increase the immunostimulatory effect of both LPS and OVA (Figure 1) potentially originated from the carbohydrate chain.

In conclusion, we have identified that SlpB from *L. brevis* JCM 1059 is a glycosylated with *N*-glycan. The glycan chain from SlpB involved in DC interaction through DC-SIGN, and in immunomodulation. Further analysis suggested that 4 fragments of SlpB from 2 carbohydrates binding motifs are glycosylated, while the remaining 3 carbohydrates binding motifs which are non-glycosylated might have involved in cell wall binding. The sequences of these fragments were identified, and all of these fragments were from N-terminal of SlpB.

## Methods

### Materials

*L. brevis* JCM 1059 was obtained from Japan Collection of Microorganisms; THP-1 cells were obtained from RIKEN Cell Bank; antibody against DC-SIGN was obtained from Santa Cruz Biotechnology (California, USA); enzyme linked immunosorbent assay (ELISA) kit for IL-12/IL-23 (p40) was obtained from R&D Systems (Minnesota, USA); ELISA kit for IL-6, IL-10 and IL-17A were obtained from Biolegend (California, USA); PNGase F was obtained from N-zyme Scientific (Pennsylvania, USA); trypsin (L-1-tosylamido-2-phenylethyl chloromethyl ketone treated) was obtained from Worthington (New Jersey, USA); DPP-IV was obtained from AT Gen (Korea); Cy5 monoreactive dye was obtained from GE healthcare (Buckinghamshire, UK). C_18_ column (5 μm, 4.6 × 150 mm; catalogue # 186003116) was obtained from Waters (Ireland); ZipTip C_18_ was obtained from Millipore (Cork, Ireland). Other reagents were obtained from Nacalai Tesque (Kyoto, Japan).

### Purification of SlpB from *L. brevis*

SlpB was purified from *L. brevis* JCM 1059 as described in (Tan et al., 2022). SlpB were stained with Cy5 monoreactive dye according manufacturer protocol as necessary.

### Adjuvant effect of SlpB

THP-1 cells were maintained and induced into DC as described in (Tan et al., 2022). To evaluate adjuvant effect of SlpB on DC, DCs were incubated with 100 ng ml^-1^ LPS with 10 μg ml^-1^ filtered sterile SlpB or 300 μg ml^-1^ OVA with 10 μg ml-1 SlpB. All reagents used are filtered sterile. SlpB was tested with limulus amebocyte lysate assay to confirm that endotoxin was below detection limit (Tan et al., 2022).

### Identification of DC-SIGN as receptor to SlpB

DC were fixed with 5%formalin solution and washed with PBS thrice. Then, DC were either incubated with antibody against DC or in PBS for 60 min at 37°C, followed by 10 μg ml^-1^ Cy5-SlpB for another 60 min at 37°C after washing with PBS. After incubation, the cells were washed thrice and analysed with fluorescent microplate reader.

### Identification of SlpB as glycoprotein

SlpB (2.5 μg) was analysed with phenol-sulphuric acid assay (DuBois et al., 1956) to identify carbohydrate contained in SlpB. Then, SlpB was co-incubated with PNGase F for 24 h according to manufacturer’s instructions to remove *N*-glycan and analysed with SDS-PAGE and stained with silver to identify *N*-glycan in SlpB.

### Competitive binding assay of SlpB

DC fixed with 5% formalin was co-incubated with co-incubated with either 100 μg ml^-1^ Glc, Gal, Man or 10 mM EDTA for 60 min at 37°C. Then, 10 μg ml^-1^ Cy5-SlpB (molar ratio of hexoses to SlpB ≈ 2,000:1) was supplemented to DC and incubated for another 60 min at 37°C. The cells were washed thrice after incubation and the fluorescence intensity was measured.

### Identification of binding domains of SlpB

100 μg SlpB was incubated with trypsin for 20 – 24 h at 37°C according to manufacturer instructions to prepare trypsinized SlpB. Trypsin incubated under the same condition was used as control.

Trypsinized SlpB (from 50 μg SlpB) or control trypsin was supplemented to DC fixed with 5% formalin and incubated for 60 min at 37°C. Trypsinized SlpB was incubated under the same condition without DC as control. Then, the supernatant was collected from DC, and the plate was washed with PBS. The flow through was also collected and mixed to the supernatant.

Supernatant of DC incubated with trypsinized SlpB, trypsin, and control trypsinized SlpB were analyzed with RP-HPLC using the following conditions: 100% solvent A for 2 min, increasing to 20% solvent B in solvent A over 15 min, increasing to 45% solvent B in solvent A over 25 min, increasing to 100% solvent B over 5 min, 100% solvent B for 2 min, and decreasing to 0% solvent B (100% solvent A) in 1 min (solvent A: 0.1% trifluoroacetic acid in deionised water; solvent B: 0.1% trifluoroacetic acid in acetonitrile). Signal intensity was measured with absorbance at 220 nm. All peaks were manually collected when necessary.

### Binding of fragments of SlpB to DC

Fragments collected from RP-HPLC were lyophilized, and reconstitute with deionised water. Then, the fragments were stained with Cy5 monoreactive dye and desalted with ZipTip C18. The desalted peptides fragments were lyophilized and reconstitute in deionised water. Binding assay and competitive binding assay was conducted as previously described, with peptide fragments obtained from 5 μg ml^-1^ SlpB.

### MALDI-ToF-MS analysis

Non-binding fragments of SlpB were lyophilized and dissolved in 0.1% trifluoroacetic acid in deionised water. Measurements were performed using an Ultraflex II TOF/TOF mass spectrometer (Bruker Daltonics GmbH, Bremen, Germany) equipped with a solid-state laser (λ = 355 nm). The sequences obtained were eliminated from the sequence of SlpB.

For analysis of binding fragments, the fragments were co-incubated with 0.5 μg DPP-IV and PNGase F and incubated for 60 min at 37°C. Mixture of DPP-IV and PNGase F were incubated under the same condition as control. Then, the peptides fragments were collected, lyophilized, reconstituted in 0.1% trifluoroacetic acid and analyzed with MALDI-ToF-MS. The sequence were predicted based on the calculated mass of trypsinized fragments of SlpB.

### Binding of affinity of fragments of SlpB to DC-SIGN

The concentration of fragment 1 – 4 from SlpB was evaluated with *O*-phthalaldehyde assay; while the concentration of SlpB was evaluated with Bradford assay. Then, the binding affinity of fragments 1 – 4 and SlpB were evaluated with Biacore X100 (GE healthcare, Uppsala, Sweeden) with sensor chip protein G (Cytiva, Uppsala, Sweeden) based on single cycle kinetics. Sensogram from blank cycles and flow path without analyte was used as control.

### Statistical analysis

Statistical analyses were performed using GraphPad Prism 9.2.0 (GraphPad Software, San Diego, CA, USA).

## Author Contributions

NY and ZLT contributed to all processes of this study.

## Reference

DuBois, Michel., Gilles, K. A., Hamilton, J. K., Rebers, P. A., & Smith, Fred. (1956). Colorimetric Method for Determination of Sugars and Related Substances. Analytical Chemistry, 28(3), 350–356. https://doi.org/10.1021/ac60111a017

Feinberg, H., Jégouzo, S. A. F., Rowntree, T. J. W., Guan, Y., Brash, M. A., Taylor, M. E., Weis, W. I., & Drickamer, K. (2013). Mechanism for Recognition of an Unusual Mycobacterial Glycolipid by the Macrophage Receptor Mincle. Journal of Biological Chemistry, 288(40), 28457–28465. https://doi.org/10.1074/jbc.M113.497149

Fina Martin, J., Palomino, M. M., Cutine, A. M., Modenutti, C. P., Fernández Do Porto, D. A., Allievi, M. C., Zanini, S. H., Mariño, K. V., Barquero, A. A., & Ruzal, S. M. (2019). Exploring lectin-like activity of the S-layer protein of Lactobacillus acidophilus ATCC 4356. Applied Microbiology and Biotechnology, 103(12), 4839–4857. https://doi.org/10.1007/s00253-019-09795-y

Geijtenbeek, T. B. H., van Vliet, S. J., Koppel, E. A., Sanchez-Hernandez, M., Vandenbroucke-Grauls, C. M. J. E., Appelmelk, B., & van Kooyk, Y. (2003). Mycobacteria Target DC-SIGN to Suppress Dendritic Cell Function. Journal of Experimental Medicine, 197(1), 7–17. https://doi.org/10.1084/jem.20021229

Guo, Y., Feinberg, H., Conroy, E., Mitchell, D. A., Alvarez, R., Blixt, O., Taylor, M. E., Weis, W. I., & Drickamer, K. (2004). Structural basis for distinct ligand-binding and targeting properties of the receptors DC-SIGN and DC-SIGNR. Nature Structural & Molecular Biology, 11(7), 591–598. https://doi.org/10.1038/nsmb784

Konstantinov, S. R., Smidt, H., de Vos, W. M., Bruijns, S. C. M., Singh, S. K., Valence, F., Molle, D., Lortal, S., Altermann, E., Klaenhammer, T. R., & van Kooyk, Y. (2008). S layer protein A of *Lactobacillus acidophilus* NCFM regulates immature dendritic cell and T cell functions. Proceedings of the National Academy of Sciences, 105(49), 19474–19479. https://doi.org/10.1073/pnas.0810305105

Koppel, E. A., Saeland, E., de Cooker, D. J. M., van Kooyk, Y., & Geijtenbeek, T. B. H. (2005). DC-SIGN specifically recognizes Streptococcus pneumoniae serotypes 3 and 14. Immunobiology, 210(2–4), 203–210. https://doi.org/10.1016/j.imbio.2005.05.014

Malamud, M., Cavallero, G. J., Casabuono, A. C., Lepenies, B., Serradell, M. de L. Á., & Couto, A. S. (2020). Immunostimulation by Lactobacillus kefiri S-layer proteins with distinct glycosylation patterns requires different lectin partners. The Journal of Biological Chemistry, 295(42), 14430–14444. https://doi.org/10.1074/jbc.RA120.013934

Mobili, P., de los Ángeles Serradell, M., Trejo, S. A., Avilés Puigvert, F. X., Abraham, A. G., & De Antoni, G. L. (2009). Heterogeneity of S-layer proteins from aggregating and non-aggregating Lactobacillus kefir strains. Antonie van Leeuwenhoek, 95(4), 363–372. https://doi.org/10.1007/s10482-009-9322-y

Mohamadzadeh, M., Olson, S., Kalina, W. V., Ruthel, G., Demmin, G. L., Warfield, K. L., Bavari, S., & Klaenhammer, T. R. (2005). Lactobacilli activate human dendritic cells that skew T cells toward T helper 1 polarization. Proceedings of the National Academy of Sciences, 102(8), 2880–2885. https://doi.org/10.1073/pnas.0500098102

Möschl, A., Schäffer, C., Sleytr, U. B., Messner, P., Christian, R., & Schulz, G. (1993). Characterization of the S-Layer Glycoproteins of Two Lactobacilli. In T. J. Beveridge & S. F. Koval (Eds.), Advances in Bacterial Paracrystalline Surface Layers (pp. 281–284). Springer US. https://doi.org/10.1007/978-1-4757-9032-0_27

Prado Acosta, M., Goyette-Desjardins, G., Scheffel, J., Dudeck, A., Ruland, J., & Lepenies, B. (2021). S-Layer From Lactobacillus brevis Modulates Antigen-Presenting Cell Functions via the Mincle-Syk-Card9 Axis. Frontiers in Immunology, 12, 602067. https://doi.org/10.3389/fimmu.2021.602067

Prado Acosta, M., Ruzal, S. M., & Cordo, S. M. (2016). S-layer proteins from Lactobacillus sp. Inhibit bacterial infection by blockage of DC-SIGN cell receptor. International Journal of Biological Macromolecules, 92, 998–1005. https://doi.org/10.1016/j.ijbiomac.2016.07.096

Smits, H. H., Engering, A., van der Kleij, D., de Jong, E. C., Schipper, K., van Capel, T. M. M., Zaat, B. A. J., Yazdanbakhsh, M., Wierenga, E. A., & van Kooyk, Y. (2005). Selective probiotic bacteria induce IL-10–producing regulatory T cells in vitro by modulating dendritic cell function through dendritic cell–specific intercellular adhesion molecule 3–grabbing nonintegrin. Journal of Allergy and Clinical Immunology, 115(6), 1260–1267. https://doi.org/10.1016/j.jaci.2005.03.036

Tan, Z. L., Kitamoto, Y., Miyanaga, K., & Yamamoto, N. (2022). *Levilactobacillus brevis* surface layer protein B promotes liposome targeting to antigen-presenting cells in Peyer’s patches. International Journal of Pharmaceutics, 622, 121896.

Valverde, P., Martínez, J. D., Cañada, F. J., Ardá, A., & Jiménez-Barbero, J. (2020). Molecular Recognition in C-Type Lectins: The Cases of DC-SIGN, Langerin, MGL, and L-Sectin. ChemBioChem, 21(21), 2999–3025. https://doi.org/10.1002/cbic.202000238

